# Post-2020 Kunming 30% target can easily protect all endemic sharks and rays in the Western Indian Ocean and more

**DOI:** 10.1101/2021.03.08.434293

**Authors:** Jessica Cheok, Rima W. Jabado, David A. Ebert, Nicholas K. Dulvy

**Affiliations:** Earth to Ocean Research Group, Department of Biological Sciences, Simon Fraser University, Burnaby, Canada; Elasmo Project, P.O. Box 29588, Dubai, United Arab Emirates; Pacific Shark Research Center, Moss Landing Marine Laboratories, Moss Landing, USA; Research Associate, South African Institute for Aquatic Biodiversity, Grahamstown, South Africa

## Abstract

Sharks and rays are possibly the most threatened Class of marine fishes and their declines can be halted if protected areas are optimised to benefit these species. We identify spatial priorities for all 63 endemic sharks and rays in the marine biodiversity hotspot, the Western Indian Ocean (WIO). Collectively, while the WIO nations currently surpass the 10% Aichi ocean protection target, this amounts to a dismal protection of only 1.57% of each species’ distribution range. We show that the entire ranges of all endemics can be achieved by protecting 11% of EEZs of WIO nations, well within reach of the new 30% of oceans by 2030 target. Regional management bodies exist, which if taken advantage of to implement shark and ray management, provide opportunities to implement more efficient management across the region. We recommend key management actions to implement and explicit incentivisation of international cooperation in the post-2020 biodiversity framework.

**Science for Society:** The past decade has seen massive growth in the establishment of marine protected areas (MPAs), driven by the Aichi biodiversity target of protecting 10% of all ocean areas. This expansion of MPAs, however, has largely occurred in areas residual to extractive uses, often coinciding with less threatened areas of lower conservation value. This coming decade will see a further push to ensure 30% of the oceans are protected by 2030. It is important to understand how existing and future MPAs should be placed to benefit threatened biodiversity. Currently this is unclear for sharks and rays, comprising a species group that is the most evolutionarily distinct vertebrate radiation in the world and also one of the most threatened. We identify both regional and national conservation priorities for expanding marine protected areas to benefit all 63 endemic sharks and rays occurring in the Western Indian Ocean region. We find that the region has already exceeded the 10% ocean protection target, but this amounts to an average of only 1.57% protection of the distribution ranges of these species. We show that protecting the top 10% priority sites will conserve almost half of the geographic range of each species yet require only 1.16% of the total EEZ – a tiny fraction of the 30% by 2030 target. We also show that regional collaboration among all nations can result in more spatially efficient conservation priorities. We recommend that the post-2020 biodiversity framework needs to explicitly incentivise regional cooperation between nations to efficiently achieve urgent targets and maximise benefits to biodiversity.

## Introduction

Two decades or more after various warnings of a rising wave of marine extinctions^1–4^, the wicked conservation challenge of protecting exploited megafauna largely remains. While there has been some success in recovery of certain megafauna species populations (e.g., right, bowhead, and gray whales)^5^, sharks and rays (class Chondrichthyes) continue to be globally threatened with a high risk of extinction^6–8^. Marine protected areas (MPAs) have become a widespread management strategy to address the overexploitation of marine resources^9^, encouraged by global biodiversity targets (under the Convention on Biological Diversity^10^) to manage 10% of coastal and marine areas within MPAs and ‘other effective area-based conservation measures’ by 2020^11^. While we have failed to meet this global target (currently 7.66% of the world’s oceans are covered by MPAs according to the World Database on Protected Areas [WDPA]^12^), the last decade has seen unprecedented growth in MPAs where approximately one-third of the growth resulted from the creation of shark sanctuaries^13,14^. The placement of these MPAs, however, is often ill-suited to conserve threatened species. For example, fewer than 12 imperilled endemic sharks and rays have some percentage (10% or more) of their distribution range in a no-take protected area^14^.

Chondrichthyans, comprised of sharks, rays, and ghost sharks, are one of the three taxonomic Classes of fishes, and are the most evolutionary distinct radiation of vertebrates^15^. These species are also one of the most threatened vertebrate groups, as assessed by the International Union for Conservation of Nature (IUCN) Red List of Threatened Species, where almost one quarter of species are threatened or predicted to be threatened (i.e., Vulnerable, VU; Endangered, EN; or Critically Endangered, CR). Further, they have the smallest fraction (<25%) of species considered as at lower risk from extinction (Least Concern; LC) of all vertebrate groups assessed^16^. The activity space or home range of fishes is strongly determined by their body size^17^. Consequently, larger-bodied species range most widely and need larger protected areas^18,19^. Existing MPAs and MPA networks are generally small in size, typically less than 5 km^2^ in extent^9,20^, providing little protective benefit for the larger wide-ranging and highly mobile species (e.g., pelagic sharks and rays). However, many shark and ray species are small-bodied with small distributions, with more than 200 endemic species occurring in the waters of a single country^21^. There is evidence that MPAs can provide effective protection for these species, since the often small sizes of implemented MPAs would cover the small extents of endemic species’ regular movements^20^. For this reason, we focus our analysis on endemic chondrichthyan species, specifically those in the Western Indian Ocean (WIO) region.

The WIO region, defined here to include sub-equatorial Africa (from the Angola-Namibia border to the Kenya-Somalia border) and the Arabian Sea and adjacent waters region (from the Kenya-Somalia border to the easternmost Sri Lanka border), harbours the second greatest biodiversity hotspot for marine fish and invertebrates^22,23^. This global hotspot is across multiple dimensions of biodiversity (e.g., species richness, evolutionary distinctiveness, endemicity^24^), including for sharks and rays^15,22,25^. In this region, coastal nations and communities are heavily dependent on marine resources, which include important shark and ray fisheries. Across the Arabian Sea and adjacent waters region, shark and ray landings from industrial and artisanal fishing support up to one fifth of the world’s chondrichthyan fisheries^26^. Similarly, across the Southwest Indian Ocean, approximately 60 million people benefit from and contribute to an annual “gross marine product” (equivalent to a country’s annual gross domestic product) of at least US$20.8 billion, where sharks are a high-value commodity^27^. This region includes the oceanic waters of Comoros, Kenya, Madagascar, Mauritius, Mozambique, Seychelles, Somalia, South Africa, Tanzania, and the Overseas Territories of France, the French Southern and Antarctic Lands, Mayotte, and Réunion. It is thus critically important to plan for the conservation of shark and ray species in this region, particularly endemics, and assess their current levels of protection within existing MPAs.

To bend the biodiversity-loss curve and reverse trends of declines^28^ over the next decade, the growth in MPAs around the world needs to maximise benefits for the most threatened species groups, notably sharks and rays. Systematic conservation planning is a widely used paradigm to identify spatial priorities and guide area-based management strategies, such as MPAs. Conservation planning is a process by which limited resources are allocated in space and time to conserve biodiversity, ecosystem services, and other valuable attributes of the natural environment, and/or address socioeconomic concerns and objectives^29^. With the numerous and likely conflicting marine conservation and development objectives across the WIO region (such as sustainable livelihoods, biodiversity conservation, and fisheries management)^30^, conservation planning can help to identify spatial priorities for endemic sharks and rays while also taking into consideration social and economic objectives^31,32^. Moreover, with the development and implementation of the post-2020 CBD Global Biodiversity Framework^33^ on the horizon, it is essential to understand what has been achieved so far and what future actions are still needed for the conservation of sharks and rays.

We identify spatial conservation priorities for all 63 endemic sharks and rays (Table S1) of the WIO (red square, inset map; Figure 1), and compare the performance of priorities identified at a regional level versus national levels across the whole region. Priorities identified at the regional level (hereafter regional priorities) emulate the scenario where all WIO countries (or donors) collaborate to achieve conservation objectives, while priorities identified at national levels (hereafter national priorities) represent the context where there is no collaboration among countries (or donors) to achieve objectives. We consider priority areas with and without considering existing MPAs in the region, to assess the contribution of existing MPAs to securing shark and ray biodiversity, and to build on the on-going protection efforts in the WIO. Areas were considered protected if they are assigned a ‘designated’ status and with any of the IUCN Protected Area categories (I-VII), according to the WDPA^12^. Our analysis identifies how existing MPAs perform for different subsets of species groupings: total species distribution range extent; IUCN Red List extinction risk category; and shark and ray body sizes. Note that species distribution ranges used to identify spatial priorities reflect known distributions at the time of analysis, which are regularly updated when new knowledge emerges. We ask five questions of the spatial priorities identified across the WIO: (1) How well does the existing MPA portfolio perform for endemic sharks and rays?; (2) Where are the priority areas that are universally important to regional- and national-level planning?; (3) Where are priority areas that are either only regionally or only nationally important?; (4) What do regional priority areas translate to for individual countries, in terms of total extent and location?; and (5) How do the most efficient priorities identified perform for different species groups?

**Figure 1.**
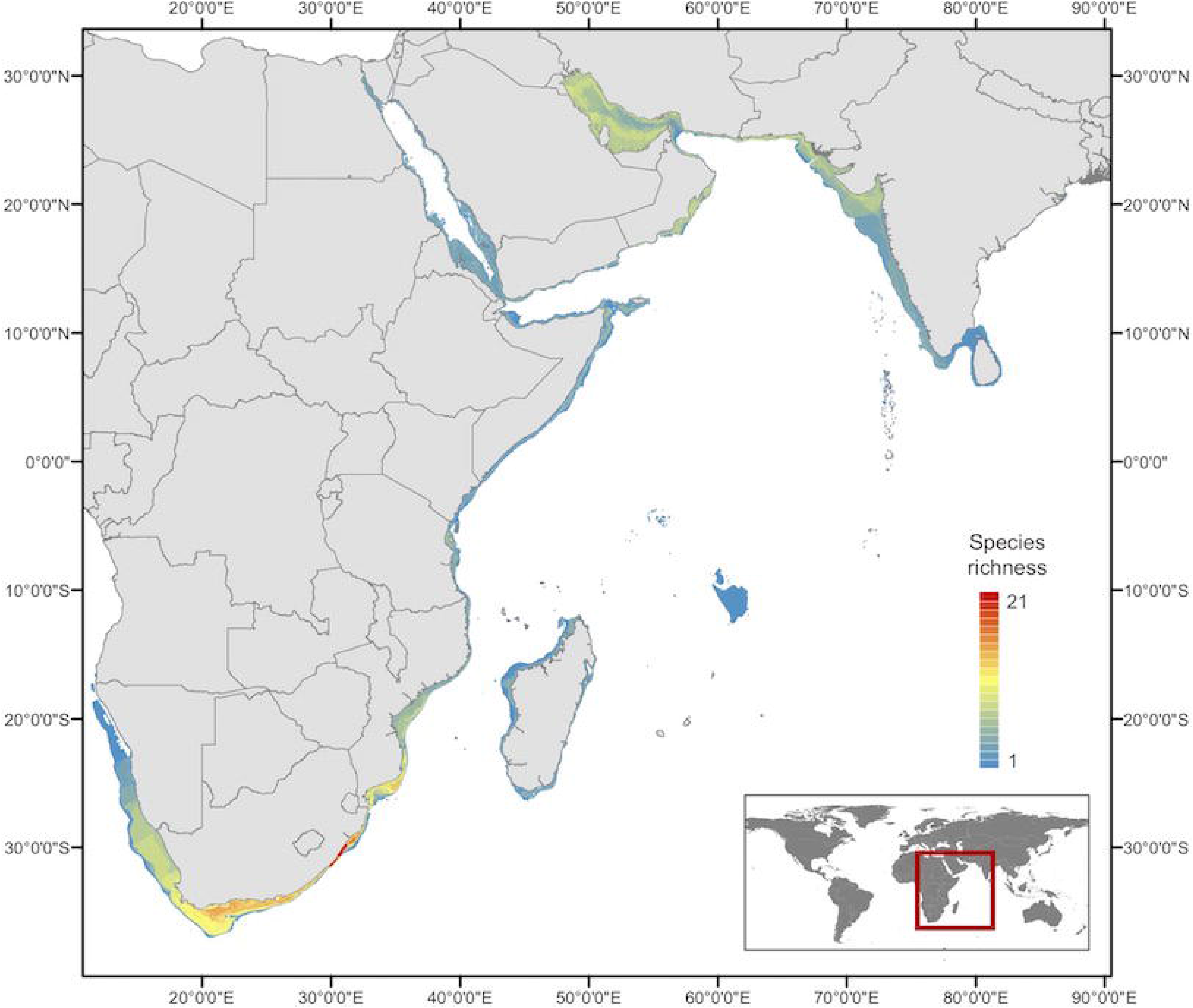
Western Indian Ocean planning region (red square, inset map) and species richness of all endemic sharks and rays in the region (*n* = 63). The Western Indian Ocean region is defined here from the westernmost point of the Angola-Namibia border, to the easternmost point of Sri Lanka’s national waters. Note that there are three endemic species with the peripheries of their distribution ranges that extend just outside of this defined region: *Acroteriobatus blochii* (Bluntnose guitarfish; extends into Angolan waters), *Triakis megalopterus* (Sharptooth houndshark; extends into Angolan waters), and *Mustelus mosis* (Arabian smooth-hound; extends into east Indian waters).

## Results

### MPA performance for sharks and rays

We find that the current MPA portfolio is not optimised to benefit sharks and rays (Figure 2). The existing MPAs in the region amount to, on average, protection of 1.57% of all 63 endemic species distribution ranges (grey shade area; Figure 2A). Moreover, there was minimal difference between priority areas identified with and without existing MPAs, and hereafter we only consider results including the current MPA portfolio. Regional priority areas were always spatially more efficient than nationally identified priorities, irrespective of whether existing MPAs were considered or not (Figure 2A). The greatest disparity between the performance of priorities of national and regional scenarios that consider MPAs, occurs at the top 24% of priority areas (B arrow; Figure 2A). Here, regional planning outperforms national planning, where there is a 10% difference in the average amount of species’ ranges protected between national and regional scenarios (67% compared to 57%; Figure 2A). Critically, when placed into the context of the total ocean area of all WIO country EEZs included in our study (31 countries; Table S2), protection of 100% of all endemic shark and ray species ranges in the region requires less than the amount of area currently under some form of protection in the region (10.76% compared to 13.5% of all EEZ areas currently protected; dark grey shade in Figure 2 inset). In other words, the 10% Aichi ocean protection target has been achieved for the WIO, but these MPAs are not located in the right places to benefit sharks and rays.

**Figure 2.**
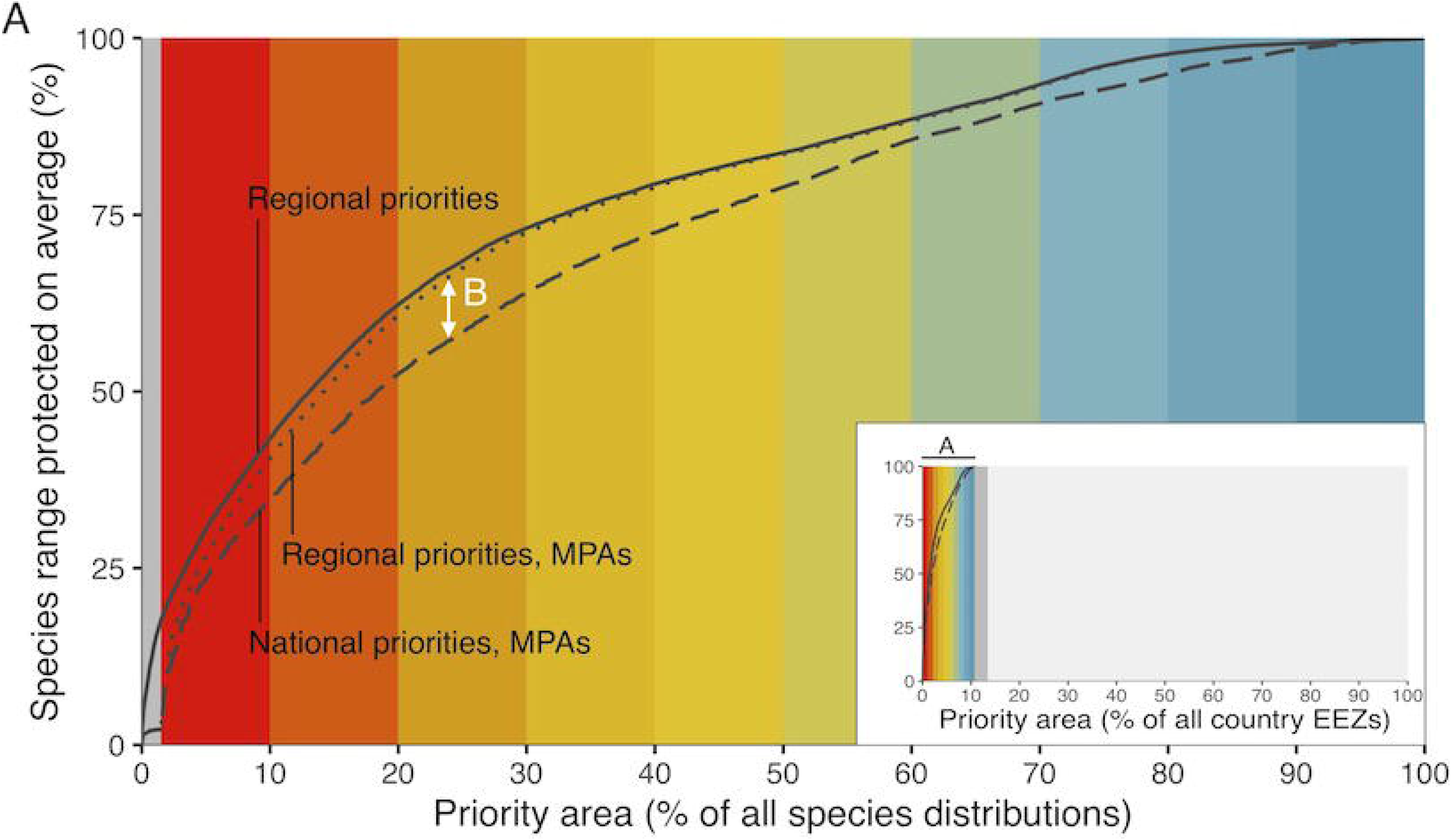
Comparison of performance of priority areas across the prioritisation scenarios. (A) Curves show the average protection of all species’ distribution ranges across the Western Indian Ocean (WIO) region for a given percentage of the planning area. The vertical grey shaded area in the main plot represents the percentage area currently under some form of protection (~1.6% of the planning region). Plot background colours correspond to the priority maps in Figure 3. Inset plot shows the same information in the context of the total area of all exclusive economic zones (EEZ) of countries (*n* = 31) within the planning area. Complete coverage (100%) of all WIO endemic species’ distribution ranges requires ~10.8% of all EEZ areas. Label A in inset plot represents the main plot.

### Universal priority areas

Some priority areas identified were universal between the regional and national scenarios (Figure 3). These areas include inshore areas off Somalia, Yemen, Oman (Figure 3c and 3d), and an area of the Laccadive Sea off southwest India (black circle; Figure 3d and 3g). In southern Africa, universal priority areas were an area off False Bay, South Africa (black cross) and the biodiversity hotspot area that stretches along the eastern South African coast into southern Mozambique coast (black circle; Figure 3e and 3h). We refined the spatial priorities identified in the regional and national scenarios to the top 10% of highest priority areas. Here, we found universally important sites in key areas across the WIO, independent of regional or national focus (Figure 4a). These universally important areas can be seen along the eastern coast of Somalia, a section of the eastern coast of Yemen, parts of the central Oman coast (Figure 4b), coastal areas around Cape Town to Port Elizabeth and coastal areas between Durban and Richards Bay in South Africa, as well areas of the southern coast of Mozambique (Figure 4c). In India and Pakistan, there are three universal priority areas: coastal areas off Hingol National Park (Pakistan), the Gulf of Kutch in northwest India, and a part of the Laccadive Sea west of the southwestern tip of India (Figure 4d). Around Madagascar, there are also three distinct areas of universal priority areas identified at both regional and national levels, at the southernmost and westernmost tips of the island, and in the north outside of Ambaro Bay (Figure 4e).

**Figure 3.**
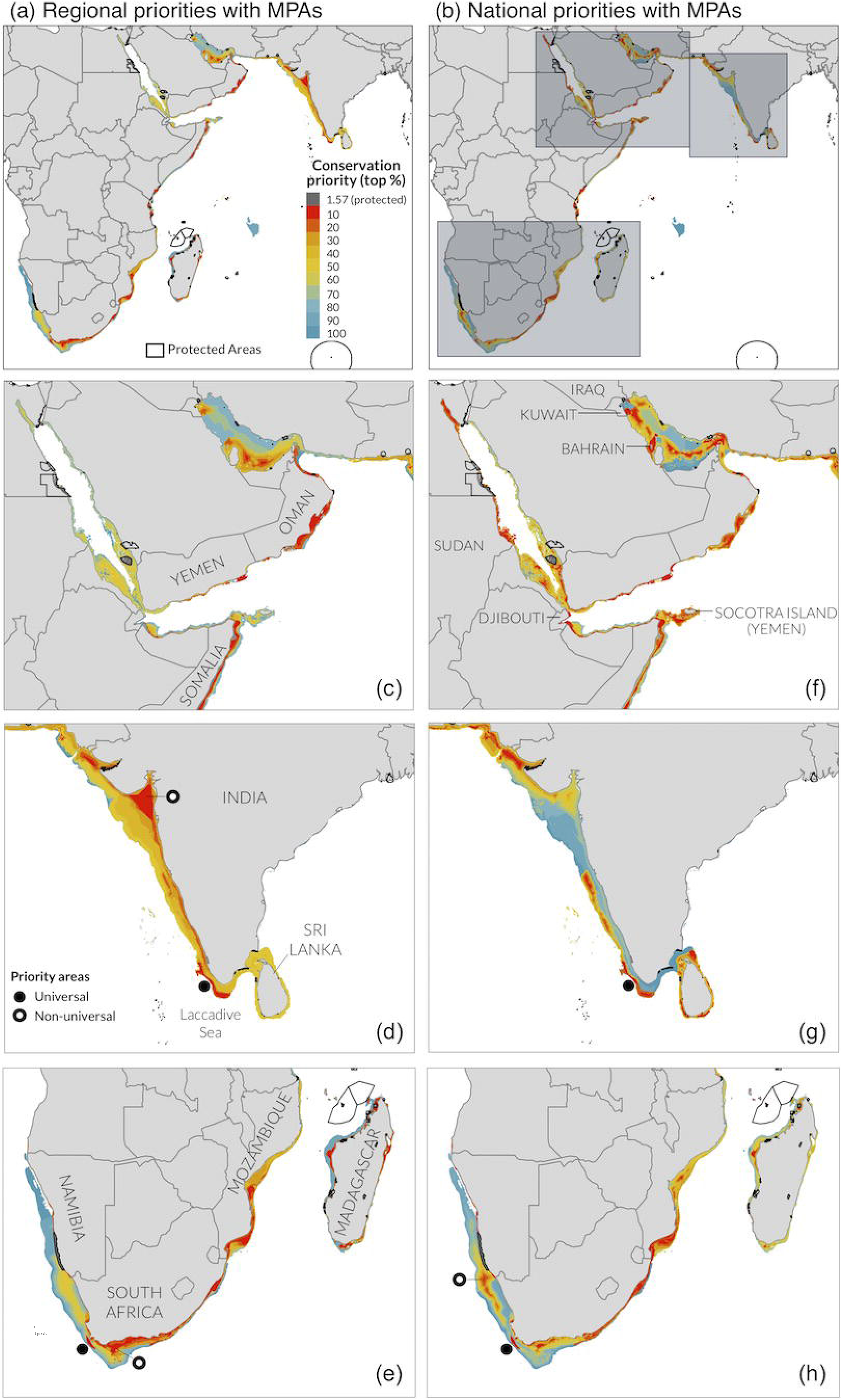
Spatial priorities across the Western Indian Ocean region between the three main prioritisation scenarios. (a) Regional priorities including existing marine protected areas (MPAs), (b) National priorities including MPAs. Inset maps (c-h) show close-up views of the main areas of differences in priorities: (c) Red Sea, Arabian/Persian Gulf, (d) India, Sri Lanka, (e) Southern Africa, Madagascar. (c-e) Regional priorities, (f-h) National priorities.

**Figure 4.**
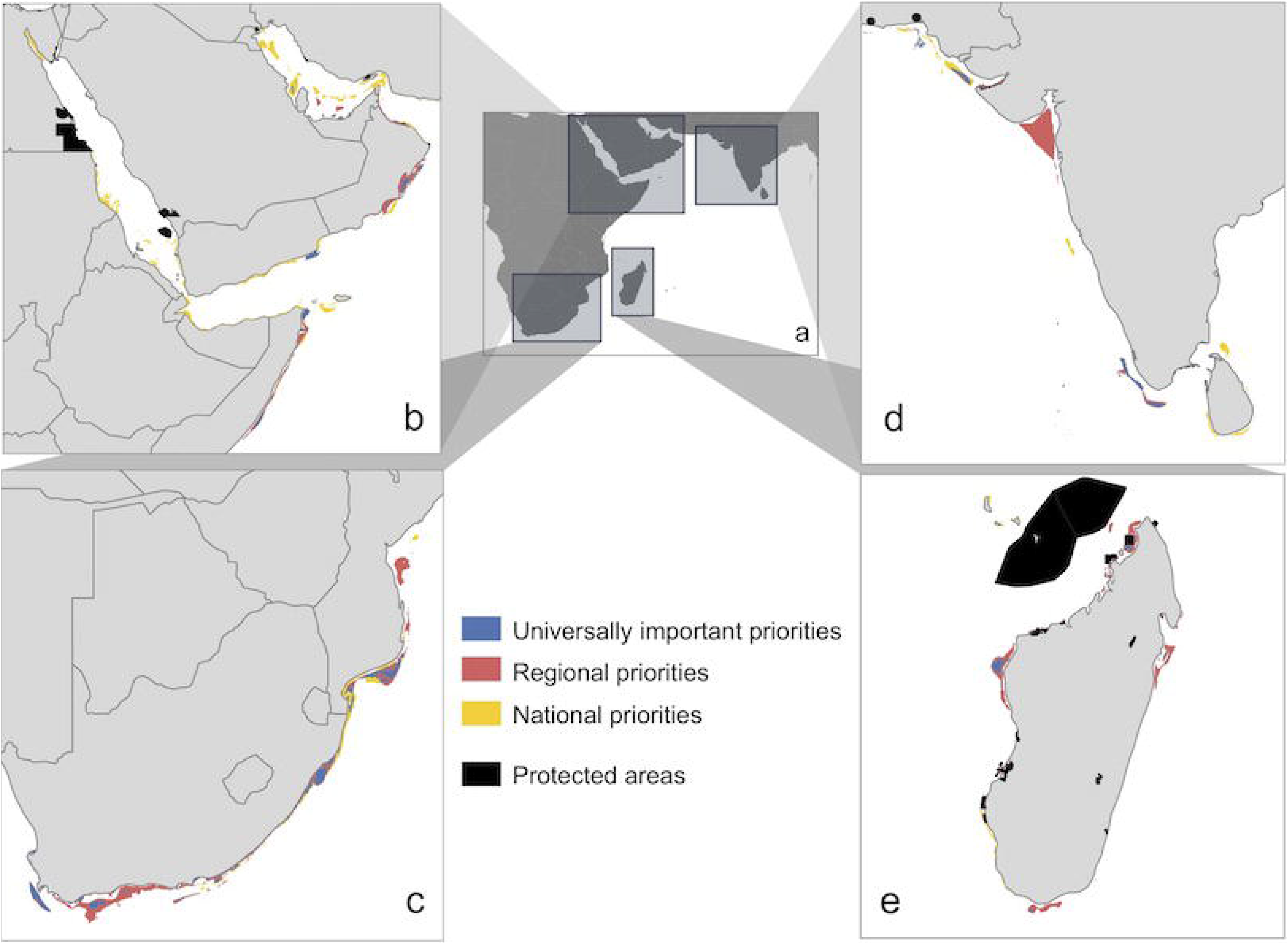
Regional and national spatial priorities (the top 10% most important areas) for all endemic sharks and rays of the Western Indian Ocean, and priority areas of overlap (~44% of top 10% of priority areas). (a) Overlapping priority areas across the whole planning region. Grey rectangles highlight regional location of inset maps, which show close-up views of the main areas of differences in priority areas (b-e). (b) Red Sea, Somalia and Socotra Island, Arabian/Persian Gulf; (c) South Africa and Mozambique; (d) Pakistan, India, Sri Lanka; (e) Madagascar.

### Non-universal priority areas

The choice of priority areas differed considerably between the regional and national prioritisation scenarios (Figure 3). Key areas of difference can be seen in the following subregions of the WIO: the Red Sea, Arabian/Persian Gulf (hereafter the ‘Gulf’), and Socotra Island; India and Sri Lanka; and South Africa, Mozambique, and Madagascar (Figure 3c-f). Across the Red Sea, Gulf, and Socotra Island (Yemen), there are a greater number of high priority areas identified in the national scenario (Figure 3c and 3f). The west coast of India has much higher regional importance compared to national priorities while conversely, coastal areas around Sri Lanka have higher national importance for the conservation and management of the endemic sharks and rays occurring in those waters (Figure 3d and 3g). Across the coastal areas of southern Africa, the main areas of differences are seen in the identified national priorities in the Orange Cone ‘Ecologically or Biologically Significant Area’ (EBSA), extending down off the western coast of South Africa (black empty circle; Figure 3h) and in the Agulhas Bank (black empty circle; Figure 3e). Around Madagascar, there are a greater number of regional priority areas identified, compared to national priorities (most notable on the northern, eastern, and southern points; Figure 3e and 3h).

### Comparison of regional and national priorities for WIO nations

The total extent of the top 10% of priority areas had a greater range between WIO nations for regional priorities than for national priorities, where the greatest extent of high-priority area within a nation’s waters was 34,769 km^2^ and 17,801 km^2^, respectively (Figure 5A and B). However, regional priorities resulted in a surprising and substantially more equitable distribution of high-priority areas between the WIO nations, ranging from 0–3.6% percent of total EEZ area, compared to 0–72.6% for national priorities (Figure 5C and D). In other words, not only are the regional priorities more efficient in achieving protection for all endemic species (Figure 2), but the resulting high-priority areas are also more evenly distributed throughout the 31 WIO national waters, relative to their total EEZ extents. In the national priorities scenario conversely, greater extents of high-priority areas were identified within each WIO nation’s EEZ due to the higher priority of nationally important endemic species. This resulted in some nations having disproportionately much greater extents of priority areas within their EEZs (e.g., Iraq, Bahrain, Kuwait, Djibouti, and Sudan; Figure 5D).

**Figure 5.**
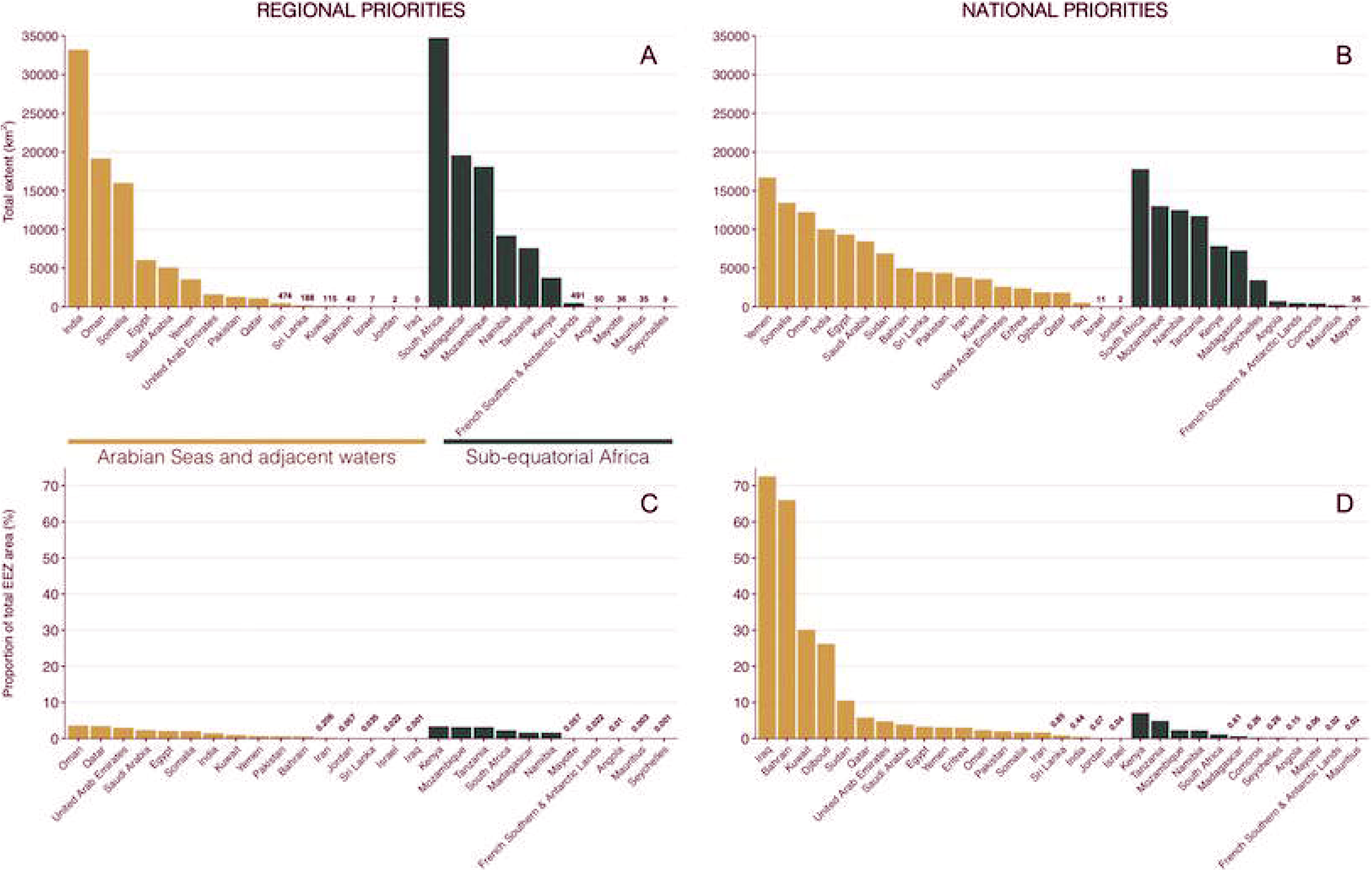
Extent of top 10% highest priority areas identified in each Western Indian Ocean nation (*n* = 31) between regional and national priorities scenarios. Comparisons are shown between regional (A, C) and national (B, D) priorities, for total extent (in km^2^) of top 10% of priority areas between nations and subregions (Arabian Sea and adjacent waters and Sub-equatorial Africa) (A, B), and as a percentage of each nation’s total exclusive economic zone area (C, D). Note the significant difference in y-axis range for proportion of total EEZ area between regional (C) and national (D) priorities.

### Performance of priorities for different species groups

We evaluated the performance of the regional priorities in terms of different groups of species according to: (1) the total extent of the species’ distribution; (2) IUCN Red List status; and (3) body size and whether the species was a shark or ray. Performances of regional priorities differed across all of the subgroups in the species (Figure 6). The total areas of species distribution ranges were similarly categorised into quartiles and as expected, regional priorities performed best in the lowest quartile, decreasing in average performance with increasing quartile (Figure 6a). In other words, the 25% smallest ranging endemic sharks and rays can have 50% of their distribution ranges protected with only 2.4% of the planning region, whereas the 25% largest ranging species can have the same amount of their ranges under protection at 45.6% of the planning region. In the context of IUCN Red List status, regional priorities performed best for Vulnerable and Data Deficient species, with approximately 50% of the distributions of species listed under these Red List categories achieving protection in 4.9% and 8.2% of the planning area, respectively (Figure 6b). Endemic species that are Near Threatened and Critically Endangered require the greatest extents to achieve 50% of species distribution ranges, ranging from 24.9% and 22.3% of the planning region on average, respectively. Size categories are different for sharks and rays (see Table S1 for size ranges assigned to different species groups). The regional priorities performed best for the one large endemic ray (Diamond Ray, *Gymnura natalensis*; located in southwestern Indian Ocean) and worst for the six medium-sized rays occurring in the WIO (Figure 6c). Approximately 50% of the distribution ranges of large rays can be protected with just 10.3% of the planning area, compared to the 55.4% of planning area needed for medium rays for the same amount (50%) of range area under protection.

**Figure 6.**
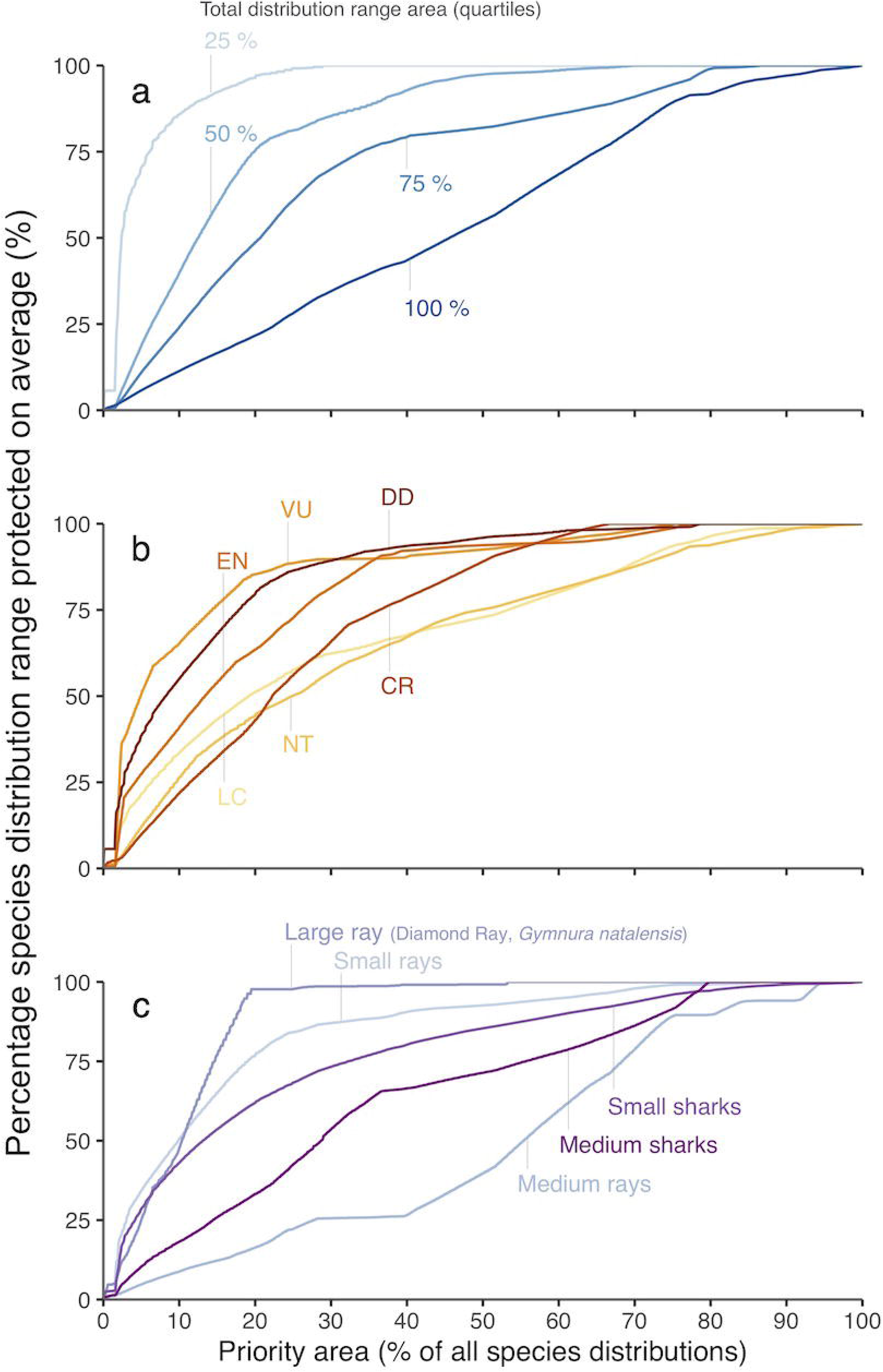
Comparison of performance of regional priority areas across different species groupings of interest. (a) Total species distribution range area categorised into quartiles (15,959 km^2^ [25%]; 58,928 km^2^ [50%]; 208,594 km^2^ [75%]; 603,085 km^2^ [100%]). (b) IUCN Red List status (Least Concern [LC]; Near Threatened [NT]; Vulnerable [VU]; Endangered [EN]; Critically Endangered [CR]; Data Deficient [DD]). (c) Body size distinguished between sharks and rays. All sharks and shark-like batoids (Rhinopristiformes): <150 cm Total Length (TL) (Small [S]); 150-250 cm TL (Medium [M]); >250 cm TL (Large [L]). Electric rays (Torpediniformes) and skates (Rajiformes): <50 cm TL (S); 50-100 cm TL (M), >100 cm TL (L). Stingrays (Myliobatoids): <50 cm Disc Width (DW) (S), 50-150 cm DW (M), >150 cm DW (L).

## Discussion

Our study demonstrates that the WIO MPA portfolio is ‘hitting the target and missing the point’^34^ in the context of endemic sharks and rays. Despite surpassing the Aichi target of 10% marine area under protection (currently 13.5% of all EEZ area is protected; Figure 2), the current MPA portfolio protects, on average, only 1.6% of all endemic species’ distribution ranges. Our findings contribute empirical evidence that achieving spatial targets alone does not necessarily ensure the protection of biodiversity^35–39^, quantifying this for the first time across MPAs in the WIO region for endemic shark and ray biodiversity. With the impending finalisation of the post-2020 Global Biodiversity Framework, there are calls for a greater spatial protection target of 30% of all land and sea^33^. It is imperative to begin implementing area-based conservation measures in places of high biodiversity value rather than residual areas of limited conservation value^40^. A recent roadmap proposes how the limitations of the Aichi target for site-based conservation can be improved by focusing on outcomes, whereby significant biodiversity sites are “documented, retained, and restored through protected areas and other effective area-based conservation measures”^41^. In line with this, we here consider and discuss (1) the extent of protection that can be achieved for all endemic sharks and rays for significant less area than is currently protected; (2) the key conservation actions to implement in priority areas; (3) how existing regional governance bodies can contribute to effective conservation; and (4) implications of spatial priorities identified for different species groups.

We identify sites of highest endemic shark and ray biodiversity value and conservation importance in the WIO and demonstrate that relatively little area is needed to protect significant proportions of their distribution ranges. If we protected the top 10% of the priority areas identified here (regional priorities, Figure 2), this would result in a 25-fold increase in the fraction of species range protected on average. Indeed, protecting this top 10% priority area would conserve nearly half (43%) of each species’ geographic range, approaching E.O. Wilson’s ‘Half Earth’ aspiration^42^, yet would take just 1.16% of total EEZ area – a tiny fraction of the 30% by 2030 target. Our calculations may be conservative because there are a number of established MPAs in the region (primarily in South Africa), currently lacking IUCN protected area category designations^12^, that could not be included in our analysis. These unclassified MPAs likely contribute some biodiversity and conservation value for endemic sharks and rays, particularly as South Africa is well-known as a regional marine biodiversity hotspot^23,25,43^. Nevertheless, empirical evidence persists that existing protected areas do a poor job for biodiversity, despite substantial recent increases in protected area coverage^44,45^. With our first-ever identified spatial priorities for all endemic shark and ray species across the WIO, we propose specific strategies to address conservation concerns for this group of species. While we have a narrow focus on these endemic sharks and rays, there is evidence that they can serve as ‘umbrella species’ conferring conservation benefits to other species, such as teleosts and crustaceans^46^.

The main conservation action to implement in these priority areas should be no-take marine areas where possible, given that the primary and ubiquitous threat to shark and ray species globally is targeted and bycatch fisheries^16,47,48^. This action is particularly critical for species that are currently overfished and with a high risk of extinction (IUCN Red List categories Vulnerable, Endangered, or Critically Endangered). With the region’s considerable dependence on marine resources for livelihoods and economic contribution^26,27,32^, however, we stress that other fisheries management activities are equally important to implement in the region. Moreover, there is evidence that sustainable take of shark or ray species can be the most appropriate management response, as complete bans or prohibition of trade (of meat, fins, or other body parts) have not typically been sufficient to reduce fishing pressure on sharks and rays^21,49,50^. The key to ensuring the long-term persistence of endemic shark and ray biodiversity in the WIO region, and indeed of any biodiversity in any region, is 100% effectively and sustainably managed and 0% overfishing and degradation^35^.

We recommend that WIO nations (and donors) focus on implementing the regional priorities identified since these are the most spatially efficient, and critically, are more equitably distributed across the 31 EEZs. We consider these priorities more equitably distributed because the proportions of EEZs identified as priorities are similar across nations, relative to total EEZ extent. This is important because otherwise the burden of protection would fall heavily on certain nations. This result of greater equitable distribution of priority areas reflects the spatial pattern of nations with larger marine estates as also having greater endemic shark and ray biodiversity. If there are endemic species of national importance, countries can also implement corresponding conservation actions to address those national objectives. Moreover, the conservation and sustainable management of shark and ray species in the WIO can be more efficiently achieved if there is greater among-country collaboration within the region^51^. The coordination of efforts among WIO countries can yield multiple benefits: saving limited conservation resources (e.g., for implementation and enforcement), improving management efficiency, and a higher return of wide-scale (i.e., regional) benefits on investment of conservation funds^52^. The Convention on the Conservation of Migratory Species of Wild Animals^53^ is the only international agreement with a Memorandum of Understanding relating to sharks and rays (Sharks MoU) with explicit objectives for enhancing regional and international cooperation^54^. Notwithstanding the focus on ‘migratory’ species, the Sharks MoU can serve as a precedent for existing regional governance bodies or agreements to explicitly encourage further collaboration between nations. While transboundary governance is rarely discussed in the context of endemic species, most of the WIO endemic species included in our analysis occur in the waters of more than one nation (50 out of the 63 endemic species are distributed across the EEZs of at least two nations).

There are existing regional fisheries and marine governance bodies that cover the breadth of our focal region, which provide a forum to encourage transboundary collaboration to manage the region’s endemic shark and ray species. While these bodies have existed for over a decade, sharks and rays have not explicitly been on their management agendas. The mandates of these bodies could be used to elevate the management status and importance of shark and ray species, particularly those endemic to the region. For the Arabian Seas and adjacent waters, there is the Regional Commission for Fisheries (RECOFI) and the Regional Organisation for the Conservation of the Environment of the Red Sea and Gulf of Aden (PERSGA). Under these regional bodies are agreements on: ensuring the state of marine resources, regulating fisheries, undertaking research and development in sustainable fisheries and the protection of marine resources (including cooperative projects between member states), developing regional capacity in all aspects of MPA planning and management, and supporting effective monitoring measures in fisheries management^55,56^. Moreover, under the RECOFI agreement, there is an article for establishing temporary, special, or standing committees and working groups to study and report on matters pertaining to the purposes of the Commission, and recommending on specific technical problems^55^ (such as the protection or sustainable management of endemic sharks and rays in the national waters of member states, particularly threatened species). For the sub-equatorial African subregion, there is the Southwest Indian Ocean Fisheries Commission (SWIOFC), which aims to improve governance through institutional arrangements that encourage cooperation among members in the context of fisheries management, and coordinate research and guide monitoring of fisheries, with an emphasis on joint activities that regard issues of regional or sub-regional importance^57^. Importantly, one of the responsibilities of the SWIOFC is to seek funds and other resources to ensure long-term operations. These governance bodies can be exploited to create transboundary MPAs and source regional conservation investments that will benefit the region’s marine resources^58^, including the conservation and management of the endemic sharks and rays of the region.

Differences in performance of regional priorities between the various groupings of the WIO endemic species highlight relatively ‘low-hanging fruit’, where the biggest protection gains can be attained for the least amount of area. Generally, ‘Small sharks’ and ‘Small rays’ species require less area for higher levels of protection (Figure 6c), which is likely attributable to these smaller-sized species having more restricted distribution ranges^15^. However, the one ‘Large ray’ species in the WIO (Diamond Ray, *Gymnura natalensis*) is the most spatially efficient to protect, because all other size groupings contain multiple endemic species, which cumulatively, amount to greater distribution extents in total. This ‘tiling’ of species distributions across the WIO largely explains the patterns of how the different species groupings perform.

We present the first prioritisation analysis of all endemic shark and ray species in the WIO, and we reveal a significant shortfall in protection based on the existing MPA portfolio in the region. This shortfall can easily be remedied well within the 30×30 target, but only if the percent area target is eschewed in favour of evaluating conservation progress. Instead, assessments of progress and impacts should focus on biodiversity (and ecosystem functioning), with emphasis on regional cooperation to improve efficiencies and minimise costs for individual nations. More generally, we highlight that the Convention on Biological Diversity biodiversity targets were designed so signatory nations can report on national targets. Yet, our approach finds that regional collaboration can produce a better conservation approach. We urge the post-2020 framework to find mechanisms and set targets to incentivise cooperation among nations to maximise biodiversity conservation outcomes.

## Supplemental information description

**Table S1. All endemic shark and ray species in the Western Indian Ocean (*n* = 63).** Species were categorised into different groups to assess performance of identified spatial priorities across three themes: (1) the total extent of the species’ distribution; (2) IUCN Red List status; and (3) body sizes, distinguishing between shark and ray species. Size measured as length types is represented as Total Length (TL) or Disc Width (DW). Threatened species (IUCN Red List category Vulnerable; VU, Endangered; EN, or Critically Endangered; CR) are highlighted in bold.

**Table S2. Countries and territories that overlap the defined Western Indian Ocean region (*n* = 31).** The Western Indian Ocean region is defined here as going from the westernmost point of the Angola-Namibia border, to the easternmost point of Sri Lanka’s national waters.

## Acknowledgements

We thank all members of the IUCN Species Survival Commission Shark Specialist Group and other experts who contributed to the data collation and species assessment, in particular, P.M. Kyne, R.A. Pollom, K. Herman, and C.M. Pollock. This project was funded by the Shark Conservation Fund, a philanthropic collaborative pooling expertise and resources to meet the threats facing the world’s sharks and rays. The Shark Conservation Fund is a project of Rockefeller Philanthropy Advisors. This work was funded as part of the Global Shark Trends Project. N.K.D. was supported by Natural Science and Engineering Research Council Discovery and Accelerator Awards and the Canada Research Chairs Program.

## Author contributions

Conceptualization, J.C., N.K.D., and R.W.J.; Data procurement and processing, J.C., N.K.D., R.W.J., and D.A.E.; Formal analysis, J.C.; Writing, J.C., N.K.D., R.W.J., and D.A.E.

