## Supplementary Material for "Post-2020 Kunming 30% target can easily protect all endemic sharks and rays in the Western Indian Ocean and more"

**Supplementary materials**

**Table S1. All endemic shark and ray species in the Western Indian Ocean (*n* = 63).** Species were categorised into different groups to assess performance of identified spatial priorities across three themes: (1) the total extent of the species’ distribution; (2) IUCN Red List status; and (3) body sizes, distinguishing between shark and ray species. Size measured as length types is represented as Total Length (TL) or Disc Width (DW). Threatened species (IUCN Red List category Vulnerable; VU, Endangered; EN, or Critically Endangered; CR) are highlighted in bold.

| **Family** | **Species name** | **Total extent of distribution range (km^2^)** | **IUCN Red List status** | **Body size category** | **Length (cm)** | **Length type** |
| --- | --- | --- | --- | --- | --- | --- |
| Carcharhinidae | Carcharhinus humani | 66 556.75 | DD | Small | 83 | TL |
|  | **Carcharhinus leiodon** | **109 008.9** | **EN** | **Medium** | **165** | **TL** |
| Dasyatidae | Brevitrygon walga | 367 061.57 | NT | Small | 32 | DW |
|  | **Maculabatis arabica** | **114 548.49** | **CR** | **Medium** | **61** | **DW** |
|  | Maculabatis randalli | 195 090.59 | LC | Medium | 61 | DW |
| Ginglymostomatidae | **Pseudoginglymostoma brevicaudatum** | **62 786.493** | **CR** | **Small** | **25** | **TL** |
| Glaucostegidae | **Glaucostegus halavi** | **419 984.17** | **CR** | **Medium** | **187** | **TL** |
| Gurgesiellidae | Cruriraja hulleyi | 222 099.28 | LC | Small | 59 | TL |
|  | Cruriraja parcomaculata | 55 970.71 | LC | Small | 43 | TL |
| Gymnuridae | Gymnura natalensis | 33 016.66 | LC | Large | 250 | DW |
| Hemiscyllidae | Chiloscyllium arabicum | 517 958.26 | NT | Small | 80 | TL |
| Heterodontidae | Heterodontus omanensis | 2 069.62 | DD | Small | 61 | TL |
|  | Heterodontus ramalheira | 86 821.19 | DD | Small | 81 | TL |
| Myliobatidae | **Aetomylaeus milvus** | **417 774.59** | **EN** | **Medium** | **123** | **DW** |
| Narcinidae | Narcine insolita | 6.09 | DD | Small | 36 | TL |
|  | Narcine oculifera | 6 649.25 | DD | Small | 35 | TL |
| Narkidae | Electrolux addisoni | 729.13 | LC | Small | 52 | TL |
|  | Heteronarce bentuviai | 583.19 | DD | Small | 19 | TL |
|  | Heteronarce garmani | 21 497.65 | NT | Small | 30 | TL |
|  | Heteronarce mollis | 28 501.01 | DD | Small | 25.5 | TL |
|  | Narke capensis | 154 680.08 | LC | Small | 38 | TL |
| Pentanchidae | Halaelurus lineatus | 73 050.65 | LC | Small | 56 | TL |
| Pentanchidae | **Halaelurus natalensis** | **84 854.11** | **VU** | **Small** | **50** | **TL** |
|  | Holohalaelurus regani | 322 238.64 | LC | Small | 69 | TL |
| Pristiophoridae | Pliotrema annae | 67.08 | DD | Small | 98 | TL |
|  | Pliotrema warreni | 149 635.29 | LC | Small | 136 | TL |
| Proscyllidae | Ctenacis fehlmanni | 10 079.72 | LC | Small | 52 | TL |
|  | Eridacnis sinuans | 19 317.91 | LC | Small | 37 | TL |
| Rajidae | Dipturus campbelli | 12 600.59 | NT | Small | 66 | TL |
|  | Dipturus pullopunctatus | 226 715.67 | LC | Medium | 130 | TL |
|  | **Leucoraja wallacei** | **242 769.02** | **VU** | **Medium** | **96** | **TL** |
|  | **Raja ocellifera** | **29 759.16** | **EN** | **Small** | **49** | **TL** |
| Rhinobatidae | **Acroteriobatus annulatus** | **23 033.67** | **VU** | **Small** | **140** | **TL** |
|  | Acroteriobatus blochii | 196.52 | LC | Small | 96 | TL |
|  | **Acroteriobatus leucospilus** | **132 934** | **EN** | **Small** | **120** | **TL** |
|  | Acroteriobatus ocellatus | 49 027.27 | DD | Small | 81 | TL |
|  | Acroteriobatus omanensis | 40 456.25 | DD | Small | 50 | TL |
|  | Acroteriobatus salalah | 58 928.16 | NT | Small | 62 | TL |
|  | **Acroteriobatus variegatus** | **273 824.4** | **CR** | **Small** | **75** | **TL** |
|  | Rhinobatos austini | 28 657.36 | DD | Small | 115 | TL |
|  | Rhinobatos holcorhynchus | 29 662.82 | DD | Small | 127 | TL |
|  | Rhinobatos punctifer | 489 768.06 | NT | Small | 90 | TL |
| Rhinochimaeridae | Neoharriotta pumila | 110 874.8 | LC | Medium | 65 | TL |
| Scyliorhinidae | Cephaloscyllium sufflans | 51 485.89 | NT | Small | 110 | TL |
|  | **Halaelurus boesemani** | **26 637.34** | **VU** | **Small** | **48** | **TL** |
|  | Halaelurus quagga | 48 314.08 | DD | Small | 37 | TL |
|  | **Haploblepharus edwardsii** | **60 851.49** | **EN** | **Small** | **64** | **TL** |
|  | **Haploblepharus fuscus** | **1 362.67** | **VU** | **Small** | **69** | **TL** |
|  | **Haploblepharus kistnasamyi** | **1 427.95** | **VU** | **Small** | **50** | **TL** |
|  | Haploblepharus pictus | 2 666.79 | LC | Small | 60 | TL |
|  | Poroderma africanum | 39 088.15 | LC | Small | 97 | TL |
|  | Poroderma pantherinum | 115 924.7 | LC | Small | 77 | TL |
|  | Scyliorhinus capensis | 298 192.01 | NT | Small | 122 | TL |
| Squalidae | Squalus acutipinnis | 7 081.1 | NT | Small | 85 | TL |
|  | Squalus bassi | 123 719.45 | LC | Small | 110 | TL |
| Squatinidae | Squatina africana | 435 617.14 | NT | Small | 122 | TL |
| Torpedinidae | Tetronarce cowleyi | 249 157.22 | LC | Medium | 113 | TL |
|  | **Torpedo adenensis** | **3 085.54** | **EN** | **Small** | **40.7** | **TL** |
|  | Torpedo fuscomaculata | 246 985.07 | DD | Small | 65 | TL |
| Triakidae | Mustelus mosis | 603 085.93 | NT | Small | 100 | TL |
|  | Mustelus palumbes | 295 046.47 | LC | Small | 113 | TL |
|  | **Scylliogaleus quecketti** | **7 247.31** | **VU** | **Small** | **102** | **TL** |
|  | Triakis megalopterus | 11 226.49 | LC | Medium | 208 | TL |

**Table S2. Countries and territories that overlap the defined Western Indian Ocean region (*n* = 31).** The Western Indian Ocean region is defined here as going from the westernmost point of the Angola-Namibia border, to the easternmost point of Sri Lanka’s national waters.

**Country / Territory**

Angola

United Arab Emirates

French Southern & Antarctic Lands

Bahrain

Comoros

Djibouti

Egypt

Eritrea

India

Iran

Iraq

Israel

Jordan

Kenya

Kuwait

Sri Lanka

Madagascar

Mozambique

Mauritius

Mayotte

Namibia

Oman

Pakistan

Qatar

Saudi Arabia

Sudan

Somalia

Seychelles

Tanzania

Yemen

South Africa
